# Design principles for perfect adaptation in biological networks with nonlinear dynamics

**DOI:** 10.1101/2022.12.05.519118

**Authors:** Priyan Bhattacharya, Karthik Raman, Arun K. Tangirala

**Affiliations:** Department of Chemical Engineering, IIT Madras, Chennai, India, Chennai, Tamil Nadu, India, Chennai, 600036, Tamil Nadu, India; Department of Biotechnology, Bhupat and Jyoti Mehta School of Biosciences, IIT Madras, Chennai, Tamil Nadu, India, Chennai, 600036, Tamil Nadu, India

**Keywords:** Network Dynamics, Non-linear dynamics, Homeostasis, Global adaptation, Biological Adaptation, Global stability, Signed Digraph

## Abstract

Establishing a mapping between the emergent biological properties and the repository of network structures has been of great relevance in systems and synthetic biology. Adaptation is one such biological property of paramount importance that promotes regulation in the presence of environmental disturbances. This paper presents a nonlinear systems theory-driven framework to identify the design principles for perfect adaptation. Based on the prior information about the network, we frame precise mathematical conditions for adaptation using nonlinear systems theory. We first deduce the mathematical conditions for perfect adaptation for constant input disturbances. Subsequently, we first translate these conditions to specific necessary structural requirements for adaptation in networks of small size and then extend to argue that there exist only two classes of architectures for a network of any size that can provide local adaptation in the entire state space, namely, incoherent feed-forward structure and negative feedback loop with buffer node. The additional positiveness constraints further narrow the admissible set of network structures. This also aids in establishing the global asymptotic stability for the steady state given a constant input disturbance. The entire method does not assume any explicit knowledge of the underlying rate kinetics, barring some minimal assumptions. Finally, we also discuss the infeasibility of the incoherent feed-forward networks (IFFLP) to provide adaptation in the presence of downstream connections. Detailed and extensive simulation studies corroborate the theoretical findings. Moreover, we propose a generic and novel algorithm based on a nonlinear systems theory to unravel the design principles for global adaptation.

## 1. Introduction

Systems biology is one of the most advanced disciplines of biology that attempts to explain any biological functionality as a response emanating from complex biological systems consisting of several constitutive units. This essentially leads to the adoption of either a conventional systems approach or a graph network formalism. The knowledge of the constitutive units (typically represented as nodes in a biochemical network) can be obtained from the traditional disciplines of biology. Further, it is widely believed from an increasing number of evidence that the network architecture for a given *phenotype* remains conserved across the organism space [1, 2]. Therefore, given a sufficient understanding of these constitutive units, finding the appropriate manner of interconnections *i. e, design principles* for important functionalities remains an exciting area of study that has attracted multi-disciplinary scholarship.

Adaptation, a ubiquitous mechanism in every living organism, is one such property that has been of sustained interest in the broader community of science. Adaptation is crucial in various processes ranging from bacterial chemotaxis to mammalian homeostasis of important metabolites. Typically, adaptation involves two subsequent steps– 1) sensing the external disturbance and 2) reverting to its pre-disturbed level. Therefore, evaluating the performance of a system vis-a-vis adaptation requires performance parameters that cater to both the aforementioned steps. For this purpose, Ma *et al* (2009) proposed two parameters, namely *sensitivity* and *precision*[1]. Sensitivity measures the quantum of the initial change in the output level post-disturbance, whereas precision is associated with the shift in steady-state output levels from pre-to post-disturbance. A perfectly adaptive system should provide infinite precision with a non-zero but finite sensitivity.

The problem of identifying adaptive network structures has attracted a lot of multidisciplinary attention ranging from computational sciences to mathematical systems theory, giving rise to broadly three different categories of approaches [3]. As the name suggests, the computational screening approach involves scanning the entire topology parameter sets. Previously, Ma *et al* (2008) studied three-protein networks and examined every possible topology–parameter combination assuming Michaelis–Menten rate kinetics. Interestingly, it was found that all the adaptive network structures contained either a negative feedback loop with buffer action or multiple forward paths from the disturbance-receiving node to the output node, with mutually opposite effects [1]. Subsequently, Qiao *et al* (2019) extended the framework to the stochastic scenario where the input disturbance was assumed to be stochastic. Apart from identifying the adaptive structures, a pair-wise correlation study between the performance parameters such as sensitivity, precision, and output signal-to-noise ratio (SNR) was carried out to obtain network structures capable of accomplishing the dual task of adaptation and noise filtering. For small-scale network structures, the correlation between the output SNR and sensitivity is negative, indicating an incompatibility between perfect adaptation and noise attenuation. However, it is possible to achieve both qualities if the dedicated adaptation and noise filtering modules are connected in series [4]. Apart from the aforementioned simulation-based techniques, optimization-based algorithms exist, which aim to find adaptive network structures by solving mixed nonlinear integer programming-based optimization techniques [5].

On the other hand, specific design strategies inspired by human-made systems have also been adopted to construct adaptive biochemical networks. Most of these design rules have been inspired by the seminal work of E. Sontag (2004) where it was argued that biological adaptation, in essence, is a disturbance rejection problem [6] from the perspective of control theory. From the celebrated internal model principle in control theory, a system capable of rejecting a step-type disturbance should contain an integrator. Inspired by this, Briat *et al* (2016) proposed an integral controller-based design, namely an antithetic integral controller (AIC) that can provide perfect adaptation for constant disturbances in a stochastic setting [7]. Subsequently, it was observed that the AIC-based design produces perfect adaptation at the cost of increased variance. To circumvent the overshoot of variance, additional negative feedback was recommended over and above the one customary to the controller module [8].

The systems-theoretic approach begins with defining a few performance parameters characteristic of biological adaptation. When mapped to the standard parameters of the underlying dynamical system (such as poles, zeros, gain *etc*.), these performance parameters give rise to a number of abstract mathematical conditions for adaptation. Further, using algebraic graph theoretic strategies, these abstract conditions are translated into structural requirements for adaptation [3, 9]. Previously, it has been shown in several works that perfect adaptation – infinite precision and non-zero finite sensitivity translates to a condition of zero-gain system in the domain of linear, time-invariant dynamical system [10, 11, 12, 13]. We previously used these systems-theoretic conditions to provide the network structures for perfect adaption for disturbance of small magnitude in three-node networks [13]. Subsequently, Araujo *et al* (2018) and Wang *et al* (2019, 2021) adopted a graph-theoretic approach to provide the structural requirements of perfect adaptation in the presence of small disturbances for a network of any size [14, 15, 16]. It has been argued that a network, irrespective of its size, must contain either an *opposer* or a *balancer* module to provide perfect adaptation in the presence of step-type deterministic disturbance of small magnitude. Further, Araujo *et al* conjectured that the opposer modules should contain at least one negative feedback loop for the purpose of stability [14]. Recently, we proved the conjecture and proposed further structural requirements to obtain the strictest necessary conditions for perfect adaptation [17]. In subsequent work, we also provided a method inspired by linear systems theory to provide a qualitative comparison of robustness to both the parametric disturbances and overall uncertainties across the structural possibilities. Additionally, we also provided a set of strong necessary conditions that constrict the search space for imperfect adaptation-capable topologies along with a relevant, sufficient condition for the same— This helped us provide structural refinements to improve the robustness of adaptation-capable networks in the presence of parametric and infinitesimal external disturbances [18].

Exhaustive screening, albeit being an extremely useful starting point remains computationally burdensome, thereby compromising on *scalability*. Further, the computational approaches require explicit knowledge about the rate laws for examining the structural possibilities. Therefore, the conclusions drawn from these studies remain largely confined to the assumed kinetics, thereby losing out on *generalizability*. Rule-based or specific design strategies circumvent both the issues but at the cost of being able to detect all the possible structural combinations for adaptation – indicating a possible loss of *exhaustivity*. On the other hand, although the recent advances in the systems-theoretic attempts to identify adaptive networks have made impressive strides vis-a-vis the scalability, generalizability, and exhaustivity, most of the approaches have assumed the input disturbance to be small enough such that the localized treatment of the underlying nonlinear system remains meaningful. Further, despite the variety of the disturbances, a few common assumptions that run through the preceding contributions in systems-theoretic approaches are– i) the input disturbance is of small magnitude, and ii) the underlying dynamical system is assumed to be relaxed prior to the step-type disturbance. The assumption of the input magnitude is small enough such that the states are not pushed into an unstable region (or beyond the domain of attraction of the stable operating point), allowing us to proceed with a linearised treatment of the problem. The second assumption translates to the requirement that the interval between two subsequent input step-type disturbances is larger than the system’s settling time. None of the aforementioned assumptions can be ensured in reality. On the other hand, the real-life network structures that have been obtained through experiments not only form a small subset of ones derived via the linearised treatment but also are capable of regulating the output for an astonishingly wide range of input disturbances[19, 20]. There have been several interesting discussions about the scale-invariance of biological networks and adaptation, being a specific scenario of scale invariance, can also be brought under the schema of these interventions[21, 22]. Although these interventions have proposed novel insights into building a nonlinear systems-driven framework for disturbance rejection, extending it to structural requirements still remains an open task.

In this work, we present a nonlinear systems theory-driven framework to identify the design principles for *global, perfect adaptation* in the presence of *deterministic* disturbance of arbitrary magnitude (GPAD). The adaptation capability in the global sense (anywhere in the state space) relaxes the assumptions mentioned above in the linearised treatment. Further, the nonlinear treatment renders the imposition of practical constraints on the system’s treatment of biological adaptation possible. For instance, since all the states are considered as the concentration of the biological species such as proteins, genes it is customary for the resulting dynamical system to be *positive* [23], a condition that cannot be accommodated in a linearised systems framework. To check the veracity of the framework, we first use the conditions inspired by nonlinear systems theory to obtain the topologies for perfect adaptation in a localised sense. Conceivably, the resultant network structures obtained for local analysis coincide with the structural predictions obtained using a linearised analysis. Hence, the proposed method is unique in literature, for it does not entirely rely on the Jacobian analysis of the system to provide the admissible topologies for adaptation in the presence of small input disturbance. Further, we utilise the conditions for positiveness and global stability to deduce the network structures that can provide perfect adaptation irrespective of the amplitude and time interval of the step-type disturbances.

The paper is organised in the following manner. Section 2 outlines the methodology that is used to arrive at precise mathematical conditions for GPAD. Subsequently, section 3 entails the application of the proposed methodology in determining the structural requirements for GPAD in biochemical networks. Finally, section 4 attempts to put all the results in perspective and tally the contributions vis-a-vis the existing literature.

## 2. Methodology

This section presents a generic framework inspired by nonlinear systems theory to deduce the conditions for adaptation. These conditions can then be applied to biological networks to determine the possible candidate network structures for adaptation.

Given a biochemical network with the concentration vector of the biochemical species denoted as **x**, the underlying dynamical system formulated from the reaction kinetics can be written as

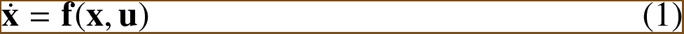

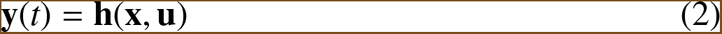

where, **u**(*t*) is the external input species of the network. We shall consider a scalar input species for the rest of this work. Further, the dynamics of the input node can be expressed as

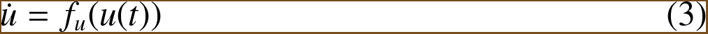

It can be seen that (1) and (3) together form a *triangular* dynamical system. According to the stability theorems on triangular systems [24], the system of the form

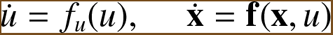

is Lyapunov stable at the origin if and only if *u*(*t*) is Lyapunov stable and **x** has autonomous asymptotically stable dynamics around the origin.

### 2.1. Assumptions

Here, we list out important assumptions on the network dynamics, which shall be valid throughout the text unless specified otherwise.

1. For an *N*-node protein system with concentrations 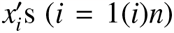 the dynamics of the concentration of *k*^*th*^ protein is assumed to be of the form 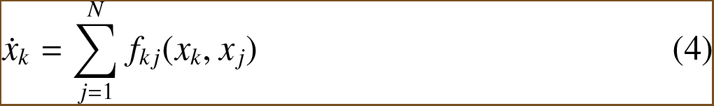
2. The flow associated with (4) satisfies semi-group property and Lipschitz continuous with respect to the states.
3. The system of differential equations constructs a well-posed system.
4. If *i* =f *j* then for a given *x*_*i*_, | *f*_*i*, *j*_(*x*_*i*_, *x* _*j*_) | is class 𝒦function with respect to *x* _*j*_ *i. e. f*_*i*, *j*_ = 0 at *x* _*j*_ = 0 and monotonic function of *x* _*j*_ in the closed interval (0, *x* _*j*_*max*).
5. |*f*_*i*, *j*_(*x*_*i*_, *x* _*j*_)| is monotonically decreasing with respect to *x*_*i*_ in the event of activation and increasing for repression.
6. In the case of activation, | *f*_*i*, *j*_(*x*_*i*_, *x* _*j*_)| = 0 when 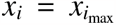 and 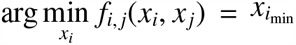 . Similarly, in the case of repression, | *f*_*i*, *j*_(*x*_*i*_, *x* _*j*_)| = 0 when 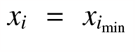 and arg max 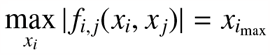
7. The input disturbance (denote it as *v*) is only connected to one protein (Denote it as *x*_1_) in an affine manner, and the effect is mediated through inter-protein connections. Further, the connection is reflected in the following way. 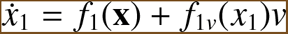

### 2.2. Conditions for perfect adaptation

In this subsection, we outline the main conditions for any biochemical network to produce the perfect adaptation with respect to bounded input with zero dynamics.

Consider a biochemical network with *N* proteins. Without any loss of generality, let us denote the concentration of the input-receiving node as *x*_1_. Further, the output is measured as the concentration of the *k*^th^ node. Given concentration vector **x**∈ℝ^*N*^ as state variables and *v* as the disturbance input, the underlying dynamics of the network can be written as

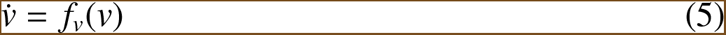

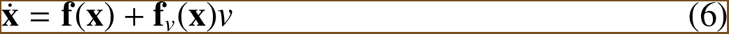

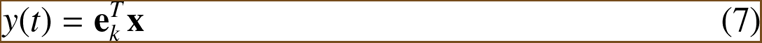

where, **e**_*k*_ ∈ℝ^*N*^ is the unit vector in the direction of *x*_*k*_ and given,

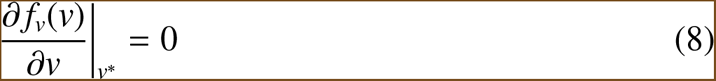

As discussed earlier, for perfect adaptation, the required non-zero, finite sensitivity can be guaranteed if the system is controllable by the disturbance input *v i. e*. it should satisfy the following condition

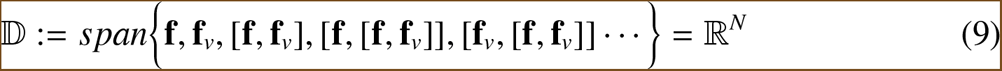

where, [**t, p**] refers to the ‘good’ Lie brackets between two vectors **t** and **p** in the tangent space of the manifold the state space evolves on.

Similarly, the infinite precision condition can be perceived as the invariance of the steady state level of the output with respect to the disturbance input. Therefore, perfect adaptation can be guaranteed if there exists a steady state (*v*^*^, **x**^*^(*v*)) in *R*^*N*+1^ such that

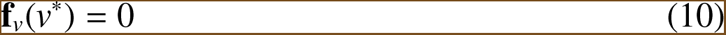

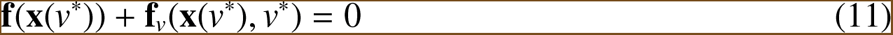

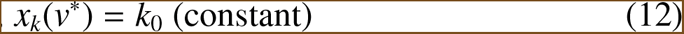

Further, the steady state must be Lyapunov stable for perfect adaptation to non-trivial initial conditions. For this purpose, we adopted a two-step procedure where in the first step, we guarantee the local stability of the system by verifying whether the Jacobian matrix at a particular steady state is stable. If yes, subsequently, we investigate whether there exists a compact, positive invariant set (Ω) containing only the concerned steady state. For this purpose, we use Nagumo’s theorem (1942) [25, 26], which provides the necessary methodology to check whether a given set is positive invariant.

#### Nagumo’s Theorem (1942)

*Assume the dynamical system in* (1) *has a unique solution for a constant u. Consider the set 𝒞*: **x**∈ℝ^*N*^, 𝒞_*i*_(**x**) ≤**b** *where, 𝒞*_*i*_(**x**) *and* **b** *are smooth functions of* **x** *such that* Δ𝒞_*i*_(*x*) ≠ ∀0**x**∈∂𝒞 *and constant column vector respectively. Further, if the set containing the active constraints is only non-empty on the boundary, then the closed set* C *is positive invariant with respect to* (1) *if and only if*

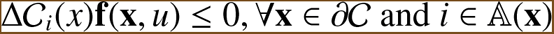

where, 𝔸 (**x**) is the set of active constraints at the boundary of 𝒞.

Finally, using a Lyapunov function (*V*) in Ω we show that for adaptation capable networks 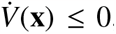, ∀**x**∈Ω, the equality holds only at 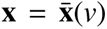, where 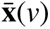 is the isolated unique steady state of (6). Therefore, from LaSalle’s invariance principle, we conclude 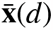 is a globally asymptotically stable steady state for a given disturbance level.

Interestingly, it can be seen that the scenario of perfect adaptation in the neighbourhood of the steady state *i. e. infinitesimal adaptation*, as termed by **(author?)** [16] satisfies the premise of the celebrated centre manifold theorem (CMT) [27]. In that case, Eq. 7-9 serves as the CMT equations, which can be formulated as the conditions for perfect infinitesimal adaptation along with the localized stability requirement.

## 3. Results

This section presents the novel insights gained by applying the proposed methodology in Figure 1 on the biochemical networks. The nodes and edges of a biochemical network refer to the biochemical species and interactions, respectively. Further, as defined in the preceding chapters, each edge in the network can be of two types– activation or repression leading to the emergence of a plethora of structural possibilities. Although the question of establishing global stability for biochemical networks has been attempted in considerable detail, the approach has been limited to either a mass-action or synergy kinetics [28] or a particular type of dynamics that assumes a radial unboundedness of the rate kinetics concerning a particular state [29]. Therefore, the present intervention attempts to provide a closure to the stability analysis by focusing explicitly on the networks with rational rate functions– these include Michaelis-Menten or Hill kinetics and their variants.

**Figure 1.**
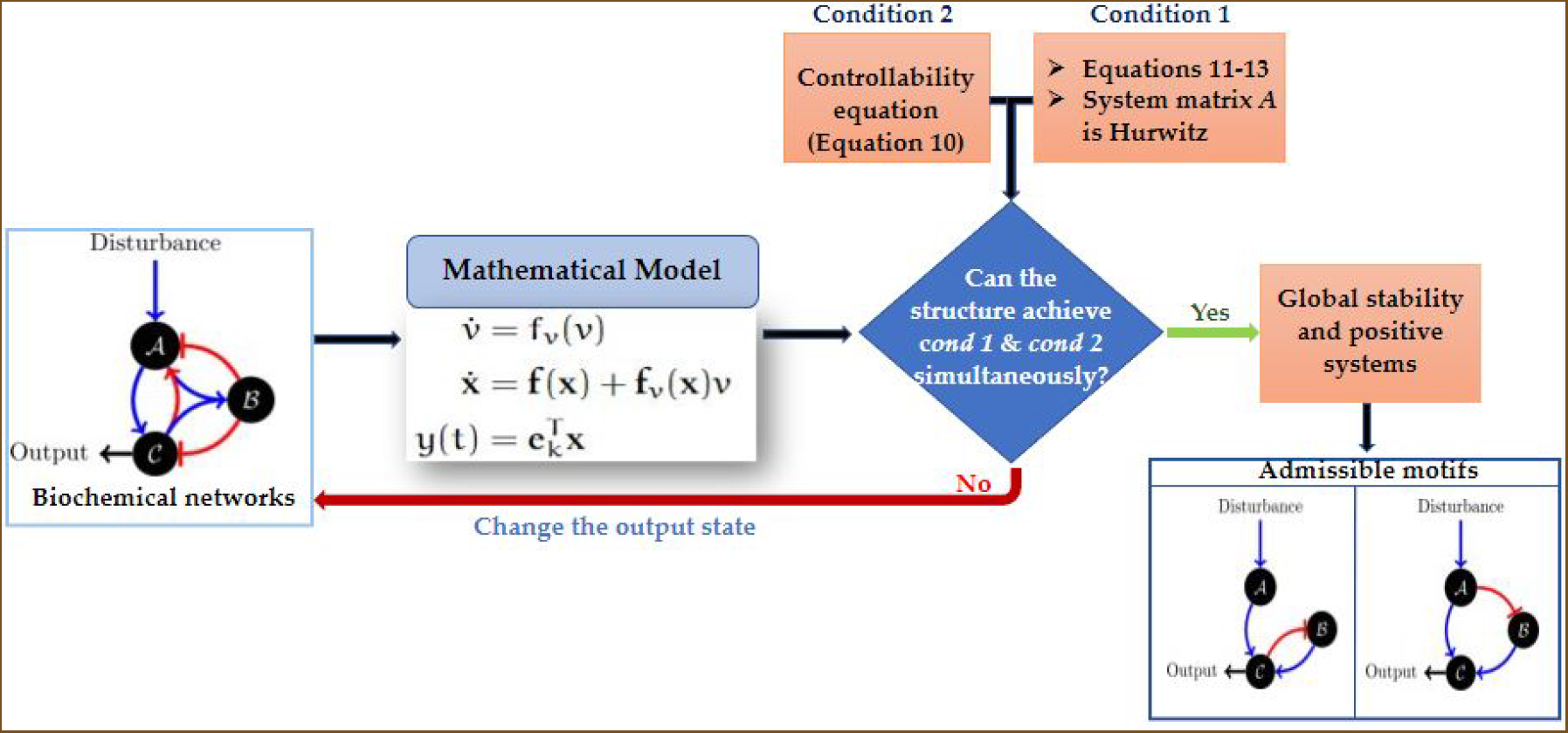
Workflow of the proposed methodology. Any given protein network is first linearized, and the conditions on the **A** matrix are investigated to derive admissible motifs for the desired functionality ultimately.

We begin with deriving network structures that can provide perfect adaptation in an input disturbance of small magnitude. Further, we assume that two subsequent step-type disturbances are further than the settling time of the dynamical system. To this purpose, we adopt a bottoms-up approach wherein we first attempt to find the minimal motifs (both in terms of nodes and edges) that can perform perfect adaptation.

### 3.1. Two node networks

As it can be inferred from the figure, adaptation is a non-monotonic response. Therefore, the possibility of achieving perfect adaptation from a single-node network (single-state system) can be safely ruled out. The immediate scenario of the two-node network can be examined.

#### Proposition 1.

*A 2-node network with the dynamics satisfying Assumptions* 1, 2, 5, *and* 7 *can perfectly adapt against constant disturbance if it contains negative feedback and a buffer action at the non-input-receiving state*.

As a corollary of Proposition 1, the two-node networks can not provide adaptation for different input receiving and output nodes— This potentially limits the application of two-node networks for in reality, despite the abundance of negative feedback loops in cell-signaling networks, the output is considered to be downstream of the external input-receiving species.

### 3.2. Three node networks

The two-node networks, as discussed before, cannot provide perfect adaptation when the input-receiving node cannot provide perfect adaptation. To circumvent this problem, we conceived a controller node (ℬ) with the following strategy

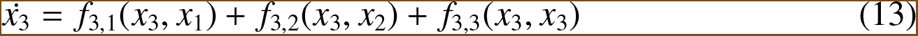

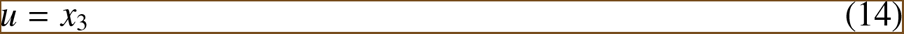

In the case of the minimal network (not more than three edges), we consider that the controller receives the information about the manipulated state through a single reaction and actuates the control signal through another chemical reaction. Therefore, they can be four such elementary network structures possible.

#### Proposition 2.

*A three-node network satisfying Assumptions* 1, 2, 5, *and* 7 *can provide local, perfect adaptation if it contains a negative feedback loop engaging the controller node acting as a buffer*.

*Proof*. Similar to the proof for Proposition 1

Fig. 2 demonstrates the response of a locally adaptive module with a negative feed-back loop with buffer action provided by the controller module. It is to be noted at this juncture that a three-node network can also have a feed-forward structure. We investigate the scenario via the following proposition.

**Figure 2.**
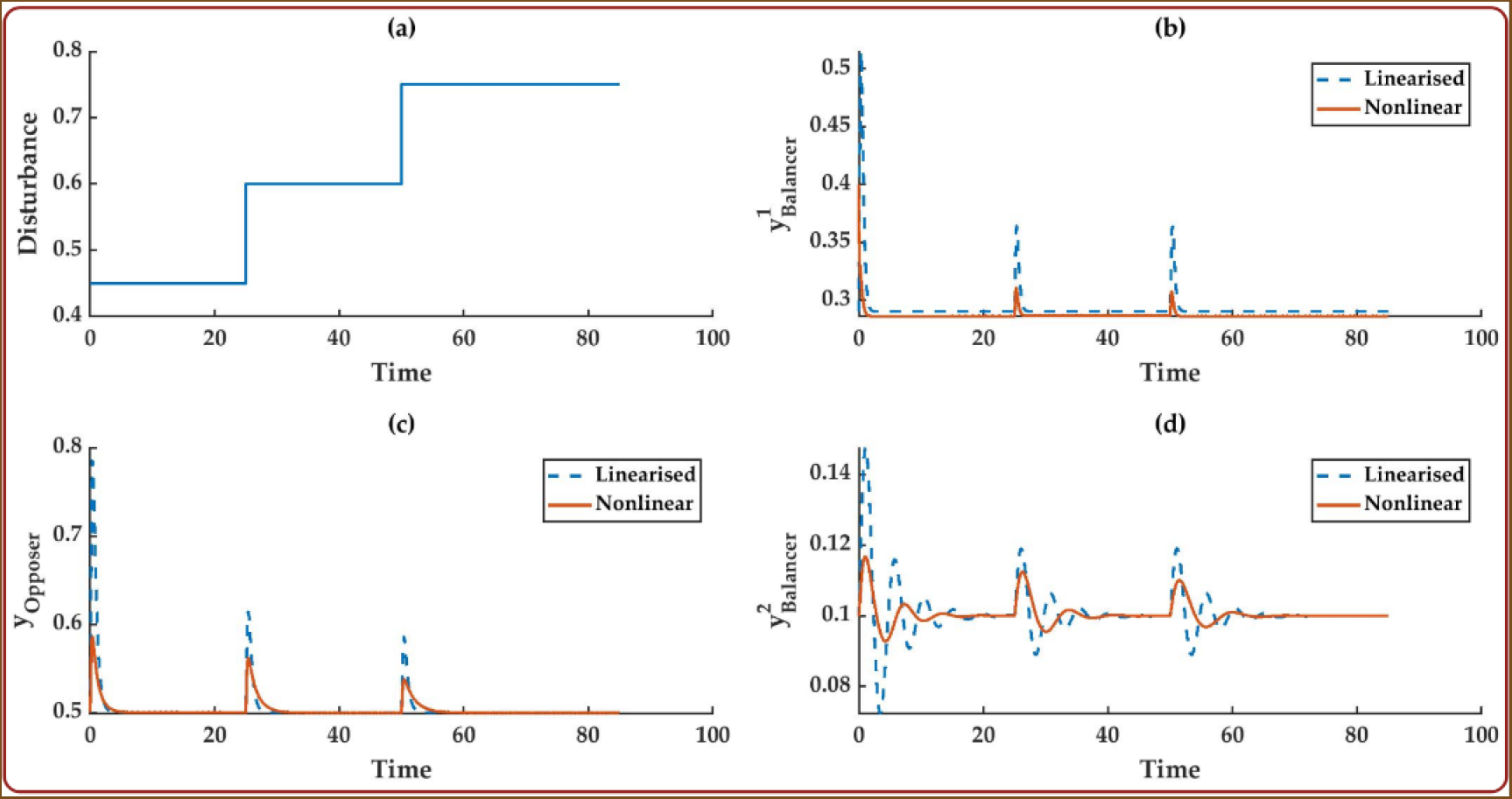
Validates the claims made in Propositions 2 and 3. The initially relaxed networks provide a perfectly adaptive response in the presence of an external input disturbance of small amplitude. Similar responses can also be obtained from the linearised analysis, which, in this case, remains valid due to small input perturbation.

#### Proposition 3.

*Three-node feed-forward network structures satisfying Assumptions* 1, 2, 5, *and* 7 *can provide local, perfect adaptation only if multiple forward paths exist from the input-receiving to the output node with mutually opposite effects*.

To demonstrate the central claim of Proposition 3, we considered three-node network feed-forward network structures with mutually opposing forward paths and Michaelis-Menten kinetics (Please refer to section 2 in the supplementary material for detailed proof). As seen from Fig. 2, the three incoherent feed-forward topology produces adaptive response against small input perturbation.

In the first three scenarios, the final network structure contains negative feedback, along with the controller dynamics being independent of the present concentration of the controller species. This class of network structures is termed as *negative feedback loop with buffer node* (NFBLB). Unlike the NFBLB, the end structures in the fourth scenario entail two mutually opposing forward paths from the input to the output node, termed the *incoherent feed-forward with proportional node* (IFFLP).

### 3.3. General structural requirements for perfect, non-infinitesimal adaptation

The analysis of two and three-node networks in the preceding subsections provides only the minimal networks acceptable for perfect adaptation in a localized sense. Since perfect adaptation, in reality, is attained by large biological networks, it necessitates a separate analysis of whether the generic design principles, such as negative feedback or incoherency in the forward paths obtained in the small-scale network analysis, are retained for networks of any size.

#### Theorem 1.

*An N*−*node network, with the underlying dynamics satisfying Assumptions* 1, 2, 5, *and* 7, *requires at least N*−*edges to provide local, perfect adaptation*.

Although Theorem 1 (Please refer to section 4 of Supplementary material for detailed proof) provides the lowest bound on the number of edges, it does not provide much insight into the network’s structural characteristics (loops, forward paths, edge types). Therefore, we further delve into the two broad structural possibilities (networks with or without loops) to determine the specific signature connections for perfect adaptation.

#### Theorem 2.

*An N-node, controllable (by the input-receiving node) feed-forward network with the underlying dynamics satisfying Assumptions* 1, 2, 5, *and* 7 *performs perfect adaptation if 1) there exist multiple forward paths from the input-receiving node to the output node, and 2) at least a pair has mutually opposite effects on the output node*.

*Proof*. Given an *N*-node feed-forward network, the external input is connected to the node *x*_1_(*t*). The maximum number of nodes *x*_1_(*t*) can influence is *N*−1. It is to be noted that for structural controllability, there exists at least one forward path from *x*_1_ to every other node in the network. Hence, no incoming edge is possible at *x*_1_ in a feed-forward network.

Let us consider the set 𝒩_1_ (*𝔎* (𝒩) = *k*_1_, where 𝔎 refers to cardinality) consists of all the nodes that contain an incoming edge from the node *x*_1_. These nodes are further connected to the downstream networks. We define the reachable set ℛ_*j*_ at a node *x* _*j*_ as the set of all nodes with an incoming edge from *x* _*j*_. Further, ℱ_*j*_ contains all the forward paths from *x*_1_ to *x* _*j*_. For instance, if a three-node network contains two forward paths from *x*_1_ to *x*_3_ such as *x*_1_ → *x*_2_ → *x*_3_ and *x*_1_ → *x*_3_ then

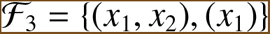

For a feed-forward network without any loops, it can be seen from Theorem 1 that nodes that have only a single incoming path from node *x*_1_ (hence the input disturbance) cannot produce the perfect adaptation. Therefore, a given node *x* _*j*_ should be considered for adaptation if

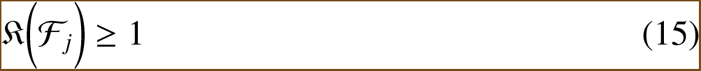

This is only possible if *x* _*j*_ is a common member of the reachable sets of at least two nodes (denote, *x*_*i*_ and *x*_*k*_) provided the ordered set (*x*_*i*_, *x* _*j*_) or (*x*_*j*_, *x*_*i*_) does not belong to any of the sets in ℱ_*j*_. The consideration of two nodes is without any loss of generality.

Suppose the node *x* _*j*_ satisfies (15) *i. e*. multiple forward paths exist from *x*_1_ to *x* _*j*_. Fur-ther, since the network is structurally controllable from *x*_1_, a break-away node (*x*_*ba*_) exists from which two paths diverge. Therefore, the dynamics of *x*_*p*_ and *x*_*q*_ can be expressed as two functions *g*_*p*_(*x*_*ba*_) and *g*_*q*_(*x*_*ba*_) respectively. Then, the adaptation equation for the output node *x*_*k*_ can be written as

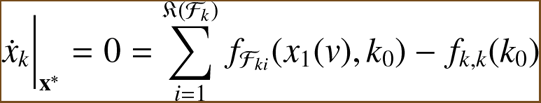

Where, 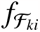 is the contribution of the *i*^th^ forward path in ℱ_*k*_ to the dynamics of *x*_*k*_, *k*_0_ is the constant steady-state concentration of *x*_*k*_. Therefore, for a step change in the disturbance

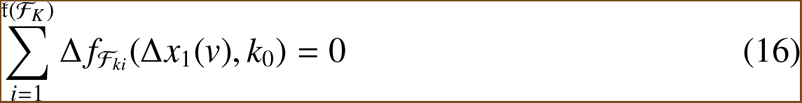

(16) can only be satisfied if there exists at least a pair of elements in 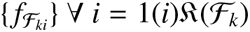 has opposite signs. Let us consider without any loss of generality that the contribution of ℱ_*p*_ and 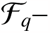 the paths consisting of nodes *x*_*p*_ and *x*_*q*_ respectively have opposing effects in the dynamics of *x*_*k*_. This can only happen if the cumulative signs of these two paths after *x*_*ba*_ are opposite, implying incoherency.

Theorem 2 provides the vital requirements for perfect adaptation in a feed-forward network. As a next step, we shall investigate the scenario of loop networks wherein we shall attempt to provide a set of structural conditions that can be safely eliminated from the set of admissible network structures for perfect adaptation.

#### Theorem 3.

*An N-node single-loop, controllable network without incoherent feed-forward paths cannot provide perfect adaptation if the loop contains an edge from the output to the input-receiving node*.

*Proof*. Consider an *N*-node network 𝒢 with a single loop (*L*_*p*_) involving *N*_*p*_ number of nodes including the input-receiving node. The controllability condition guarantees at least one path from *x*_1_ to every other node in the network. Suppose, there exists an edge from any node *x*_*l*_∈*N*(𝒢) *N*(*L*_*p*_) (where *N*(𝒢) returns the node set of the network) to *x*_*p*_∈*N*(*L*_*p*_). It is to be noted here since *x*_*p*_ ∈ *N*(*L*_*p*_), there exists a path from *x*_*p*_ to *x*_1_. This implies the existence of a loop other than *L*_*p*_ involving the forward path from *x*_1_ to *x*_*l*_ and *x*_*l*_ to *x*_*p*_ and *x*_*p*_ to *x*_1_– this violates the condition of a single loop.

Since there exists only one loop and no upstream connection from the non-loop nodes to the nodes pertaining to *L*_*p*_, it is possible to show through constructive proof (Supporting information) that the resulting dynamics of the *L*_*p*_ can be expressed as

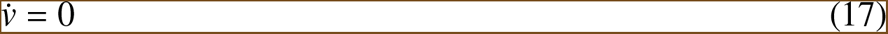

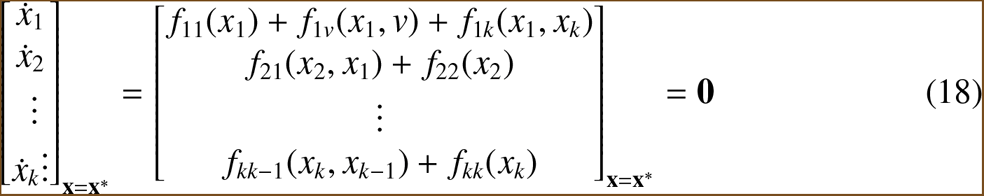

Putting the infinite precision condition 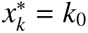 on equation 18 and applying the assumption that 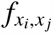 is a class-K function of *x* _*j*_.

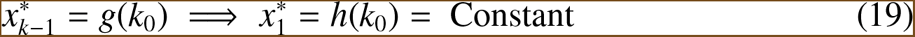

It can be observed that for (18) and (19) to be consistent if and only if the external disturbance level *v* is kept constant, which defies the entire purpose of adaptation. Therefore, this has to be treated as a contradiction.

It is to be noted that, although Theorem 3 suggests the presence of only one loop, the main result works even if multiple loops are involving only those nodes that do not figure in *L*_*p*_. Further, even in the case of multiple loops with an edge from the output to the input node, it can be shown that the network structure fails to provide perfect adaptation (Supporting information).

From Theorem 1, it is only appropriate to consider networks with at least *N* edges. Similarly, Theorem 3 also eliminates the set of network structures where all the loops include an edge from the output to the input-receiving node. Further, Theorem 2 excludes the coherent feed-forward networks from the set of admissible topologies for perfect adaptation. Therefore, we shall only investigate those network structures that pass the check-points proposed in Theorems 1, 2, and 3.

#### Remark 1.

*As demonstrated in the foregoing sections, for an N-node network to provide perfect adaptation, it has to provide infinite precision* ((11) *and* (12)*) along with non-zero sensitivity (*(9)*). It is to be observed that* (11) *is a system of N equations. From our analysis of two and three-node networks, we have seen a particular equation in the system of equations in* (11) *that guarantees the constant steady-state value of the output state. We shall denote this equation as the* ***invariance equation***. *Suppose there exists a particular node x*_*c*_ *in the network such that the following relation holds*

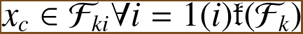

*where ℱ*_*k*_ *contains all the forward paths from the input-receiving node to the output node. Further, x*_*c*_ *is the immediate node before x*_*p*_ *for every forward path in ℱ*_*k*_. *In this scenario, it can be shown that x*_*p*_ *inherits the response of x*_*c*_ *in the context of adaptation, provided there is no loop between x*_*c*_ *and x*_*k*_. *Therefore, the structural requirements that guarantee perfect adaptation for x*_*c*_ *are sufficient for x*_*p*_.

#### Remark 2.

*Further, the invariance equation can be either at the output node x*_*k*_ *(or x*_*c*_ *if it exists) or any node other than x*_*k*_. *As the next step, we shall investigate both these two cases separately*.

*In the first scenario, the dynamics of the output node or x*_*c*_ *(if it exists) can be written as*

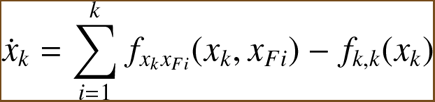

*where, k refers to the incoming degree of x*_*k*_ *and x*_*Fi*_ *is the immediate node before x*_*k*_ *for the forward path ℱ*_*ki*_.

*According to the assumption*, 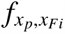 *are class 𝒦 function of x*_*Fi*_, *the possibility of making it independent of x*_*Fi*_ *can be safely eliminated. Therefore, the only way the equation corresponding* 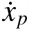 *can become the invariant equation is if, for a change from v*_1_ *to v* + Δ*v, the following condition holds true*

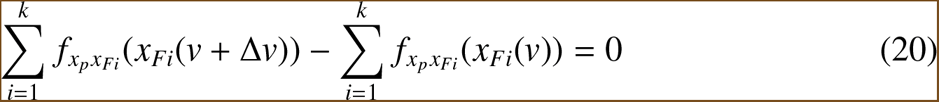

*It can be proved that due to the class 𝒦 nature of f*_*i*, *j*_(·) *the steady state solution* **x**^*^(*v*) *to* (6) *is a monotone function of v. Therefore*, 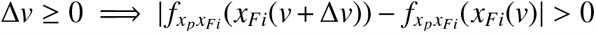. *Therefore* (20) *can only be satisfied if at least one forward path exists whose effect on the dynamics of x*_*k*_ *is opposite to that of the rest of the forward paths. It is to be observed that none of the following conclusions change when x*_*c*_ *exists. Instead of focusing on the equation concerning* 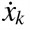, *the dynamics of x*_*c*_ *should be considered*.

#### Remark 3.

*In the second case, we examine the scenario where the invariance equation is neither the output state equation nor the one corresponding to x*_*c*_. *We assume that the invariance equation is situated at the state equation of a node x*_*b*_ *(x*_*b*_ ≠ *x*_*c*_, *x*_*b*_ ≠ *x*_*k*_*). It is evident that since the invariance equation ensures the constant steady-state value of the output state x*_*k*_ *has a path from the output node x*_*k*_. *Therefore, the dynamics equation for x*_*b*_ *can be written as*

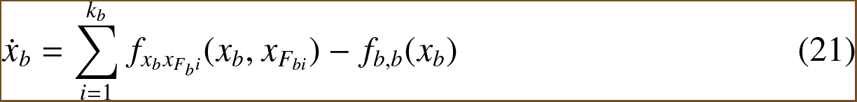

*It is to be noted that at least one of* {*F*_*b*_*i*} *has a path from the output node x*_*k*_. *Considering the class* 𝒦 *nature of* 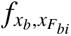, *the only way* (21) *serves as the invariance equation at the steady state rendering* 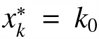 *is if* 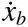 *becomes independent of x*_*b*_. *In this manner, the second term of* (21) *remains a constant, and the first term is independent of x*_*b*_, *thereby making x*_*k*_ = *k*_0_ *a possibility*.

*Further, subsequent analysis of the local stability of the system reveals that the node x*_*b*_ *has to engage in a negative feedback loop for stability (supporting information). The reason, intuitively, can be expressed in the following way. Since, f*_*x*_*b is independent of x*_*b*_, *the corresponding 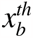 row in the system matrix* (**A**) *of the linearised system has zero diagonal component. Therefore, from combinatorial matrix theory, the elements of the* 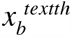 *row have to figure in the determinant expression within at least one loop expression [17]*.

### 3.4. Towards global stability

Remarks 1-3 establish the fact the structural recommendations obtained by Theorems Theorems 2, and 3 serve as necessary structural conditions for local, perfect adaptation in a network of any size. Since, by definition, global stability implies local (linearized) stability around the steady state, the structural conditions hitherto derived serve as the superset for the structures capable of global, perfect adaptation. Therefore, we begin the search for admissible network structures within the structural possibilities of incoherent fee-forward structure and negative feedback with buffer action.

It is to be noted that the states considered in the entire formalism refer to the concentration of the biochemical species (*eg*. genes or proteins). Therefore, the resulting dynamics underlying the biochemical networks constitute a positive system.

#### 3.4.1. Global adaptation for IFF

##### Claim 1.

*An incoherent feed-forward topology, controllable by the input-receiving node with dynamics satisfying Assumptions (a)-(g) provides perfect adaptation for arbitrarily large input disturbance if each node has at least one incoming activation link*.

*Proof*. Let us first define the biologically feasible range of the steady state Ω⊂ℝ^*N*+^ := **x**_min_ ≤**x** ≤**x**_max_. We shall approach the proof in two steps: at first, we prove the uniqueness of the steady state in Ω and further prove the stability of that steady state to establish the stability of the steady state for any initial condition in Ω.

#### Uniqueness of the steady state

Without any generality loss, the input-receiving node’s concentration is represented by the state *x*_1_. Given a feed-forward structure (with no cycle) with the network being structurally controllable by the input-receiving node, the dynamics can be expressed as a triangular system

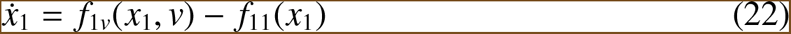

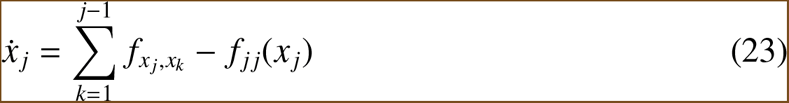

where, the rate function *f*_*ii*_(*x*_*i*_) refers to the self-degradation rate of each species. At the steady state,

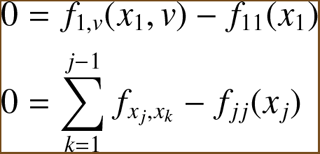

It is to be noted here that according to the statement of the claim, the rate function *f*_1,*v*_ describes an activation reaction of *x*_1_ by the disturbance input *v*. From Assumption f, for a given disturbance level *v*, |*f*_1,*v*_| and *f*_11_ are monotonically decreasing and increasing functions of *x*_1_ respectively. Further, at a non-zero positive disturbance level, 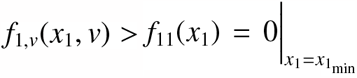 and 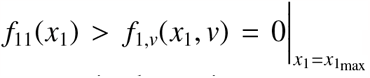 . This guarantees the existence of single, isolated steady-state 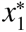 in the region [*x*_1_min, *x*_1_max]. Similarly, the other states also attain a unique, isolated solution at the steady state. This concludes the first part of the proof.

#### Stability

As it can be seen from Equations 22 and 23, that at 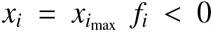, ∀**x** ∈ Ω implying 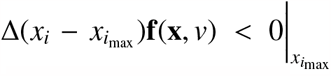 due to the customary presence of self-degradation activities and optional repression at each node. Further, due to the presence of at least one activati1ng effect present at each node 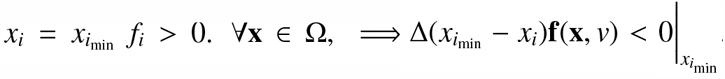. Therefore, from Nagummo’s theorem on positive invariance, we conclude that the compact set Ω is positive invariant in the current setting of feed-forward networks. Further, the system’s Jacobian obtained around the only steady state **x**^*^∈Ω adopts a lower triangular structure due to the feed-forward network structure’s tri-angular structure. Due to the customary presence of the self-degradation and Assumptions a, f, and g, the diagonals of the Jacobian are always negative anywhere in Ω rendering the only steady state **x**^*^∈Ω locally stable. Now, since Ω is the biggest positive invariant set for the dynamical system containing a single singularity **x**^*^, which, in turn, is locally stable, all the trajectories starting at Ω have to converge to **x**^*^ for the well-posed dynamical system making the system asymptotically stable in Ω. This concludes the proof. Fig. 3 demonstrates this result via a five-node balancer module in the presence of disturbance undergoing large amplitude swings. Fig. 4 lays out different feed-forward structures and elucidates this theoretical result.

**Figure 3.**
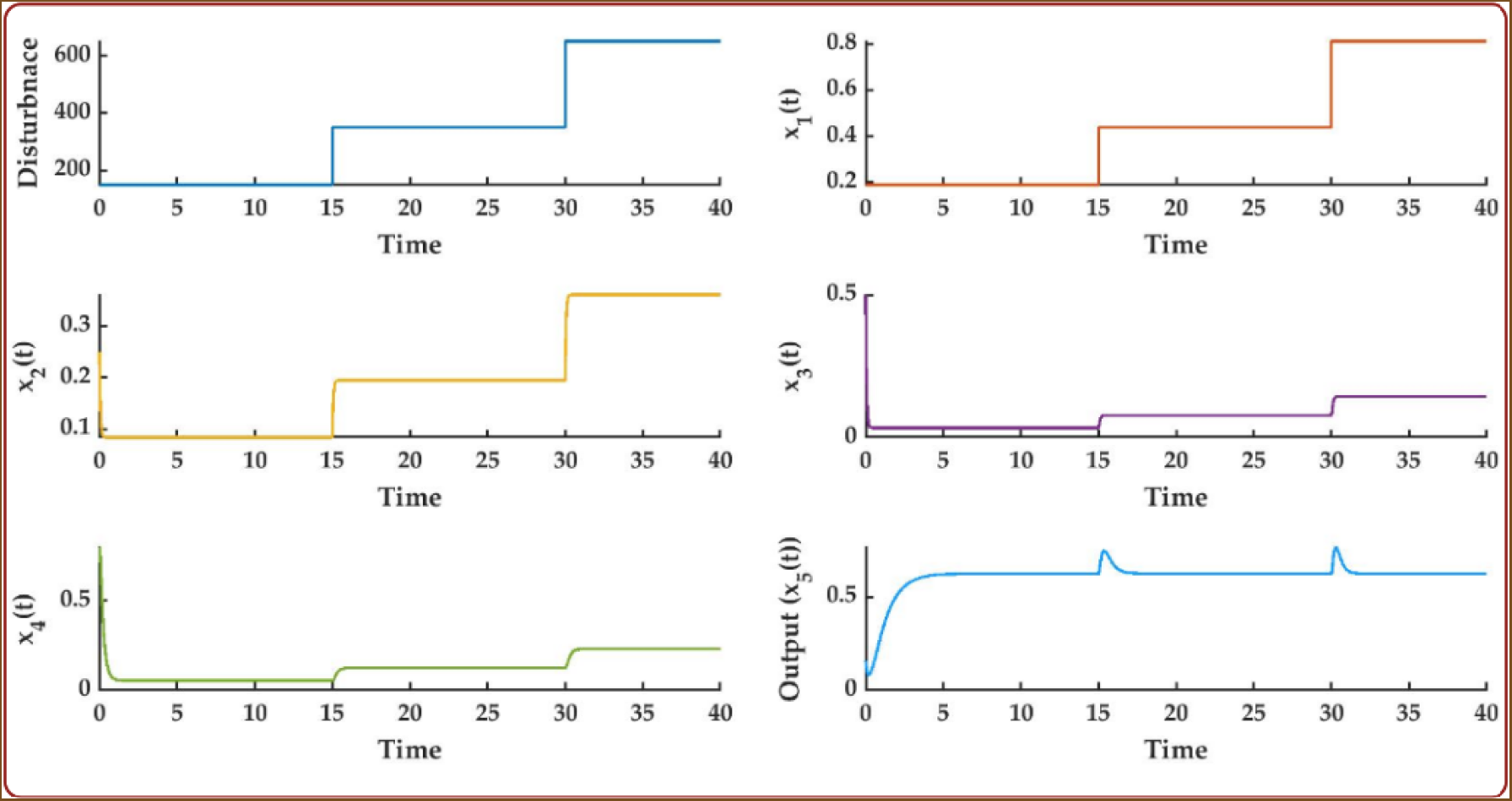
Encapsulates the conclusion drawn from Claim 1. Although the input disturbance is moved from an initial value of 0.3 to 600. The five-node balancer module retains its adaptive property. Further, as depicted in the claim, all the states lie within Ω := **0** ≤**x** ≤**1**. The initial stiffness of the output state is due to the non-steady state initial condition.

**Figure 4.**
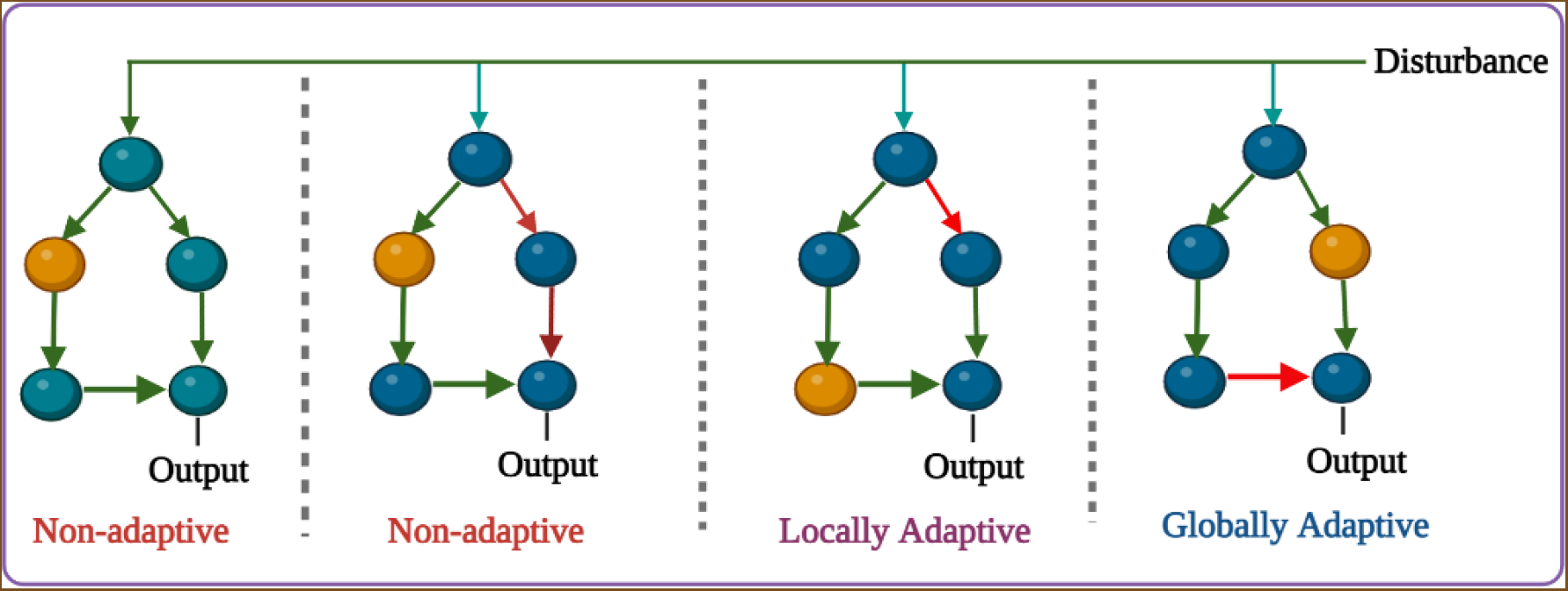
Different feed-forward structures relevant for perfect adaptation. The first two structures from the left fail to satisfy the opposing action, thereby failing to provide adaptation. On the other hand, the intermediate network structures, albeit satisfying the structural condition for local, perfect adaptation, cannot guarantee a unique and single steady state, thereby failing to guarantee global properties. On the other hand, the right-most network structure guarantees a unique, stable, steady state in Ω,, thereby exhibiting global adaptive properties. The node in Brown represents the controller node. The edges in Green and Red refer to activation and repression, respectively.

### 3.4.2. Global adaptation for NFB

#### Claim 2

*An N*−*node network structure containing a single feedback loop with an odd number of repressive edges and the underlying open-loop dynamics satisfying Assumptions (a)-(g) provides perfect adaptation for arbitrary large disturbance input if the following conditions are satisfied*

1. *Each node contains at least one incoming activation link*.
2. *The control action is exerted directly on the input-receiving node*.
3. 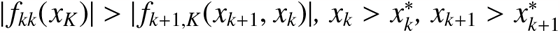 *and* 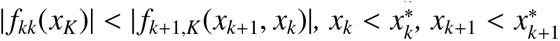

*Proof*. Similar to IFF, we also attempt to establish the claim in two steps.

#### Uniqueness of the steady state

Without any loss of generality, i) let us denote the input-receiving node concentration as *x*_1_(*t*) and ii) assume the loop engages all the *N* nodes. Since the controller action is meditated through *x*_1_(*t*), the concentration of the controller node is indexed as *x*_*N*_(*t*). Therefore, the entire network can be understood as *x*_1_(*t*) → *x*_2_(*t*) → *x*_3_(*t*) → … → *x*_*N*_(*t*) → *x*_1_(*t*). The associated dynamics can be expressed as

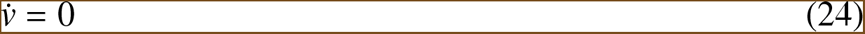

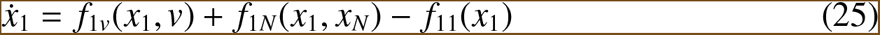

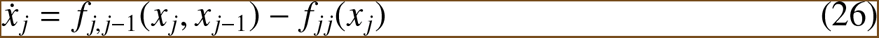

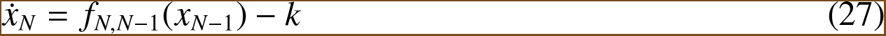

Since it is established in Theorem 3 and Remark 3 that for negative feedback network structure, the controller dynamics have to be independent of the concentration of the same Equation (27) contains no term containing *x*_*N*_ in its right-hand side. At steady state, since *f*_*N*,*N*−1_ is a monotonically increasing function of *x*_*N*−1_, the solution to Equation (27) is unique. Further, due to the customary presence of at least one incoming, activating edge at each node, the unique solution 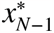 renders a unique steady state solution **x**^*^∈ℝ^*N*+^ through the preceding equations. This satisfies the uniqueness criterion.

#### Stability

The dynamical system in Equations (24) – (27) assumes a triangular form concerning (*v*, **x**). From Vidyasagar’s theorem on triangular system [30], the Lyapunov stability of the interconnected system can be established if we can establish the asymptotic stability of the autonomous dynamical system described in Equations (25) – (27). For this purpose, we propose the following Lyapunov functional

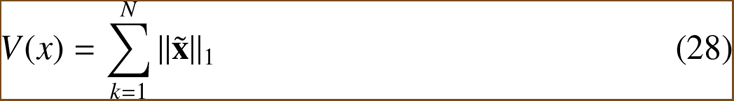

where, 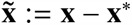. Further, since there exists a state-dependent degradation at every node from *x*_2_ to *x*_*N*−1_, the corresponding contribution of the dynamics (*f*_2_ – *f*_*N*−1_) to the time derivative 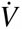 contains at least one negative sign. Further, in the special scenario of 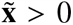 and 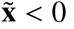, the contribution of *f*_*N*_ to 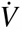 is positive. However, since *f*_1*N*_ is a repressing edge, the positive contribution of *f*_*N*_ is nullified by the same of *f*_1_ rendering the overall expres-sion 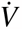 negative (it is to be noted that 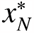 in the case of an autonomous system is unique but negative) owing to the third condition in the statement of the claim. This proves the Lyapunov stability of the interconnected system. Fig. 5 illustrates this claim through a five-node opposer module with an assumption of Michaelis-Menten kinetics. Fig. 6 provides the schematic of different possible adaptive opposer modules. Further, a simulation study for an opposer module in the presence of large disturbances is demonstrated in Fig. 5.

**Figure 5.**
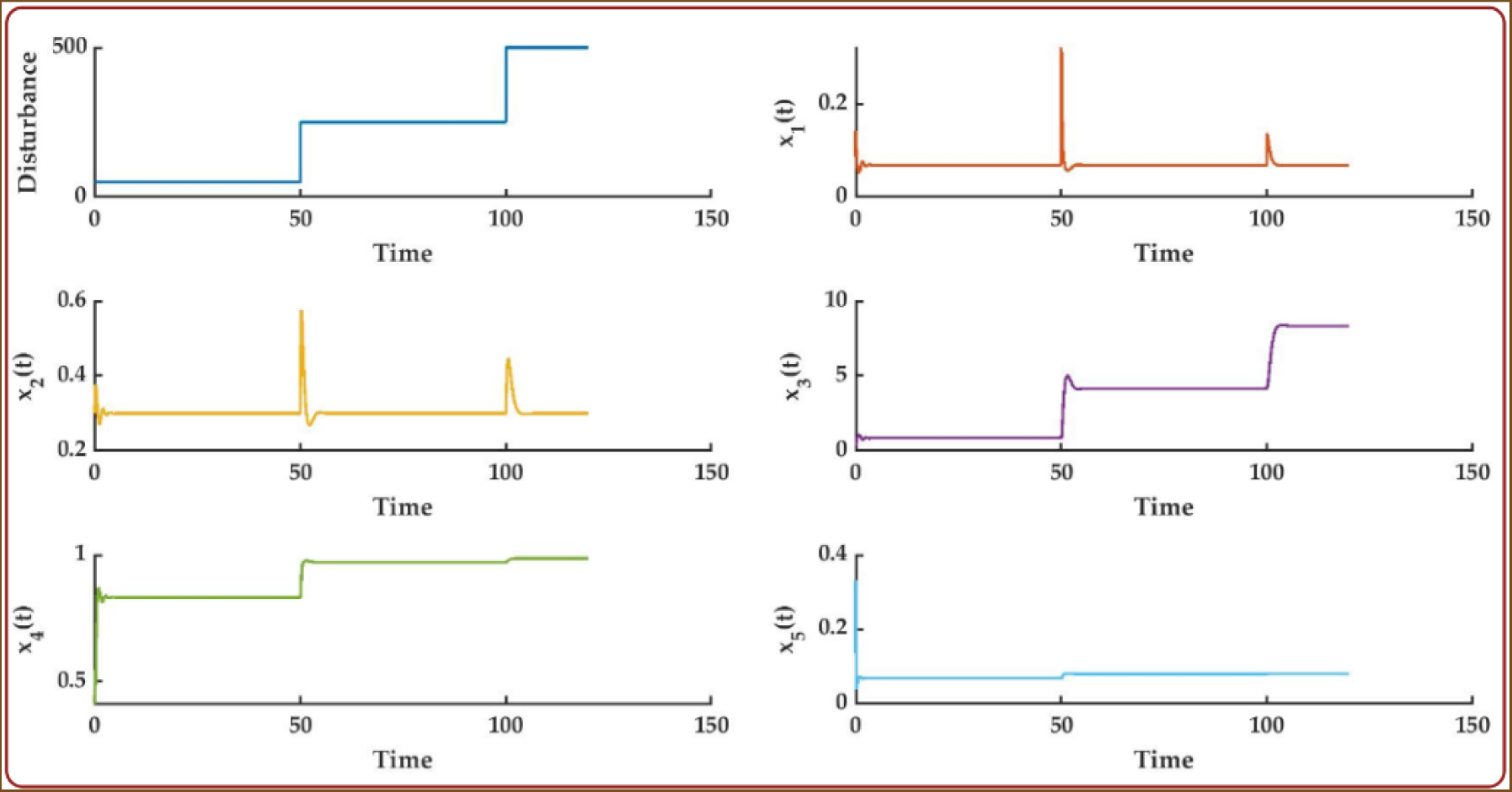
Encapsulates the conclusion drawn from Claim 2. Although the input disturbance is moved from an initial value of 0.3 to 600. The five-node, opposer module retains its adaptive property. Further, as depicted in the claim, all the states lie within Ω := **0** ≤**x** ≤**1** except the controller node. Further, it can be shown that, unlike the balancer modules, the concentration of the controller node, the node that accomplishes the buffering action, cannot be contained within [0, 1]. As it can be observed, both *x*_1_ and *x*_2_ can potentially serve as the output node of the module. Evidently, the initial stiffness of the output state is due to the non-steady state initial condition.

**Figure 6.**
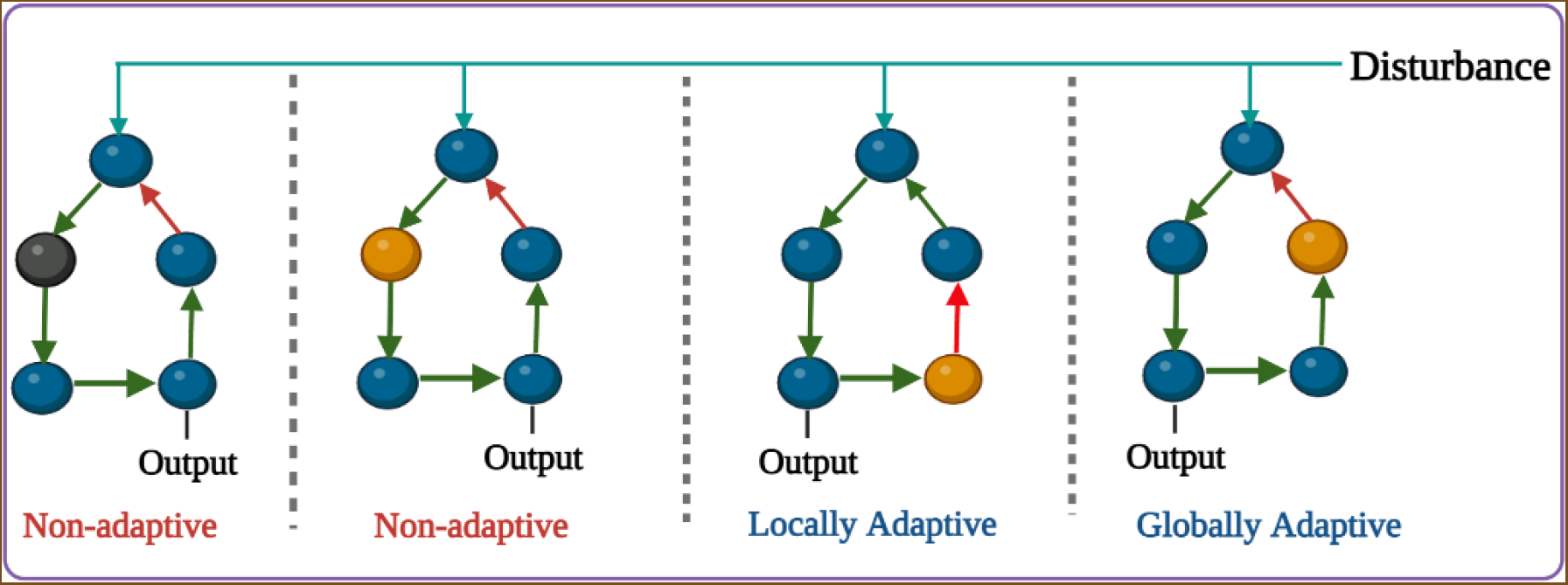
Different feed-forward structures relevant for perfect adaptation. The first two structures from the left fail to satisfy the buffer action, thereby failing to provide adaptation. On the other hand, the intermediate network structures, albeit satisfying the structural condition for local, perfect adaptation, cannot guarantee a unique and single steady state, thereby failing to guarantee global properties. On the other hand, the right-most network structure guarantees a unique, stable, steady state in the state space, thereby exhibiting global adaptive properties. The node in Brown represents the controller node providing the buffer action crucial for adaptation. The edges in Green and Red refer to activation and repression, respectively.

##### Remark 4.

*It can be seen that due to the buffer action by the controller node and Condition (b) of Claim 2, all the N*−1 *nodes of the loop (excluding the controller node) perform perfect adaptation. In general, for any loop engaging P number of nodes, P*−1 *nodes can perform perfect adaptation if the controller node exerts the control action through the input-receiving node. We can further relax the first requirement of an edge from the controller to the input node as for an P node feedback loop with the controller being the K*^*th*^ *node (K* ≤*P), the first K*−1 *nodes starting from the input-receiving node to the one before the controller node can perform perfect adaptation*.

#### 3.5. Global adaptation in the presence of downstream connection

Most of the analysis in the foregoing sections assumes that the network structure is isolated from any downstream network, which is not the case in reality. Typically, adaptation networks are mounted on top of big biological networks to improve the cell’s robustness against variations in the external environment. It is, therefore, necessary to investigate the performance of an adaptive network in the presence of downstream connections. For this purpose, we adopt the standard model of downstream connection known as retroactivity developed by [31, 32] wherein the output node of the adaptive network is connected cyclically with any of the downstream nodes. From Remark 3, it is well-known that the invariance equation is situated at the controller node in the case of negative feedback. The steady-state equation about the controller node itself decides the steady-state value of the output node. Therefore, the steady-state value of the output node is preserved against the presence of a downstream connection at the output node. This implies the stability of the network structure is not altered; the perfect adaptation property of the output node of a negative feedback network with a buffer node is conserved in the presence of a downstream network(Section 4.1 of supplementary material). Fig. 7 illustrates the modular nature of opposer modules with a simulation exercise.

**Figure 7.**
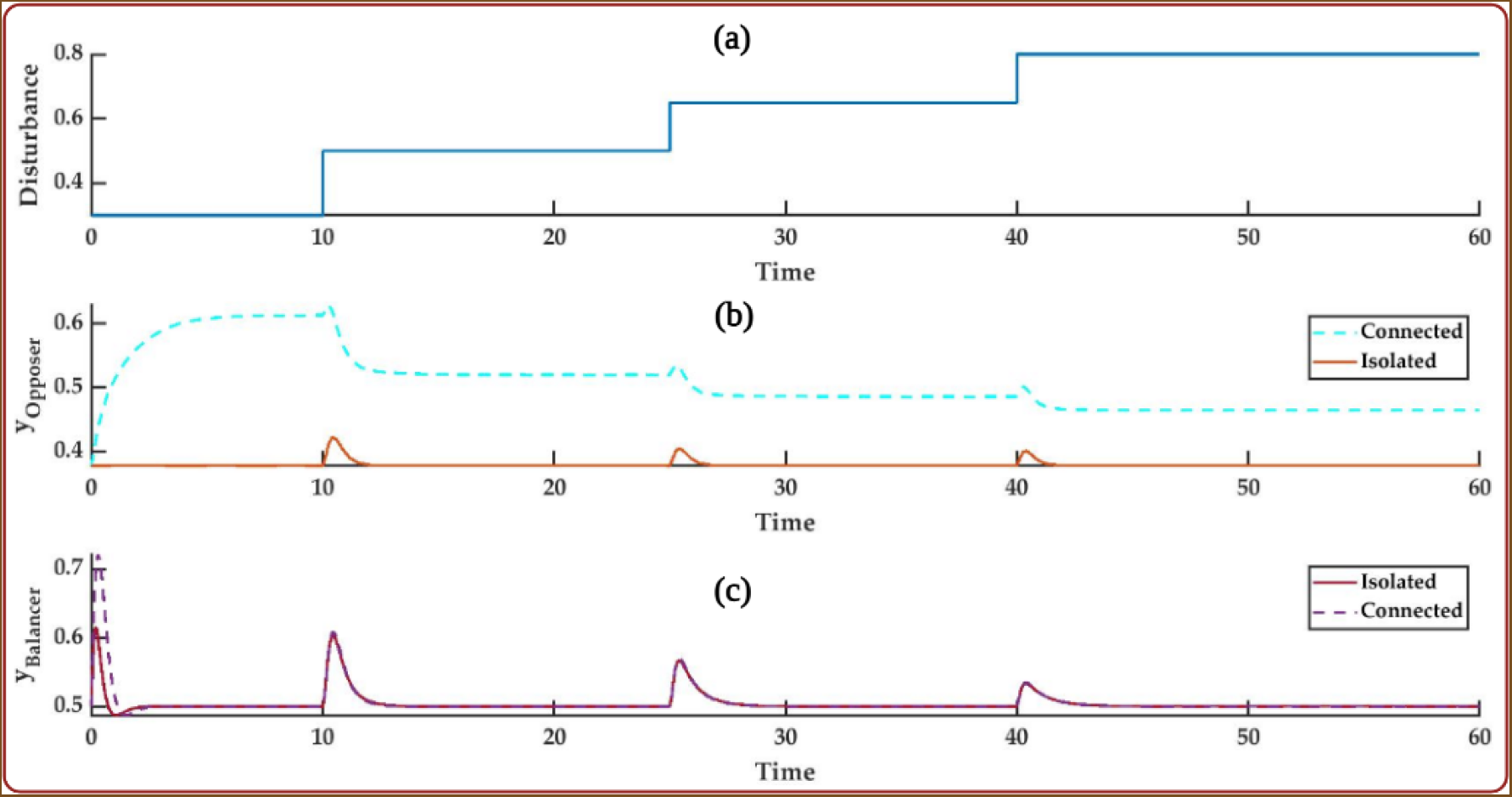
A comparative study of the retroactivity property of the opposer and the balancer modules. As it can be seen in (b), a perfectly adaptive 3−node balancer module loses its adaptation capability when connected to a downstream system consisting of two nodes. On the other hand, as shown in (c), a 3−node opposer module retains its adaptation capability even in the presence of a two-node downstream connection. Interestingly, the output response remains the same irrespective of the manner of the connections (positive or negative loops) between the output node and the downstream system.

On the other hand, the incoherent feed-forward topology, as established in Remark 2, achieves perfect adaptation through the invariance equation situated at the output node (or to another node *x*_*c*_ in the network given the output node contains nothing other than a single incoming path from node *x*_*c*_). Therefore, if the output node itself is connected to the downstream network, the invariance equation is modified. Additionally, if the downstream connection comprises feedback, the output node can oscillate around the origin, leading to a ‘negative’ value, thus an infeasible concentration value (Section 4.2 of supplementary material). Interestingly, similar results have been observed with the computational work by [33] in their study of how incoherent feed-forward loops can perform perfect adaptation (maintain constant output levels) for a specific range of retroactive strength. Fig. 7 vividly illustrates the retroactive nature of the balancer modules with a simulation exercise.

## 4. Discussion

The inherent nonlinearity and variety of possible rate dynamics contribute a major share to the complexity of the biological networks. Apart from the apparent abundance of rate dynamics, it is well-established in the literature [34, 1] that the network structure plays a governing role in determining the response of a network structure. The present study follows from this seminal observation and attempts to synthesize a methodology that provides novel structural insights without explicitly relying on the particular rate kinetics.

The existing literature, including [1, 4, 5] has focused chiefly on the computational approaches to deduce the design principles for perfect adaptation despite the obvious scalability issue. Further, rule-based approaches such as [7, 8] propose a negative feedback-based structure that is able to provide perfect adaptation. Apart from these two approaches, other theoretical interventions primarily rest on the treatment of the dynamical system in the linearised domain [16, 13, 17, 14]. This, albeit being a sophisticated approach, cannot portray the global picture– nor can it incorporate realistic constraints that can be thought of as the vital design criteria from the vantage point of the synthetic design. To circumvent this problem, the current chapter proposed a methodology inspired by nonlinear systems theory that strives to draw novel structural insights about biological networks that can produce perfect adaptation in a global sense. At first, using the classical performance parameters for adaptation, namely sensitivity and precision, we determined the conditions for perfect adaptation. Contrary to a Jacobian-based approach that is prevalent in the existing systems-theoretic approaches, we proposed the condition for infinite precision as the existence of an error-zeroing manifold in the state space. On the other hand, the sensitivity condition is met through a Lie-controllability test of the dynamical system. At this point, global adaptation is indeed a stronger requirement than an infinitesimal adaptation for the added conditions of strict global stability and a unique steady state.

The proposed framework has been used to study the biochemical networks. To verify the correctness of the algorithm, we first applied this to deduce the optimal network structures for perfect adaptation in a localised sense. Unsurprisingly, in this scenario, the structural predictions obtained are identical to the ones obtained through the Jacobian treatment of the linearised system. The proposed methodology can also deal with the case of singular Jacobians using the principle of centre Manifold Theory. In that case, instead of the concentration of a particular node, the linear combination of the node concentrations is likely to be able to produce an adaptive response.

The network structures obtained for local adaptation serve as the superset for the structures capable of global, perfect adaptation. Propositions 1, 2, and 3 provide the necessary structural requirements for local, perfect adaptation in networks of small size. Subsequently, the scope of these results is generalized using Theorems 1, 2, and 3 to establish the fact that negative feedback with buffer node or multiple feed-forward structures with mutually opposite effects on the output node is the key to local, perfect adaptation in a network of any size.

Intuitively, the structural requirements for local, perfect adaptation can only serve as the necessary conditions for global adaptive characteristics. Claim 1 establishes that ‘*not all incoherent feed-forward network can provide global adaptation*.’ Only those feed-forward structures wherein each node contains at least one activating incoming edge can provide global (in Ω), perfect adaptation. Similarly, Claim 2 also produces such a subclass of the networks containing a negative feedback loop with buffer action in the context of global, perfect adaptation. It is to be noted at this juncture that the rate law assumptions (Assumptions (6), (7)) are only used in proving the global stability of the steady state. The monotone nature of the rate functions (standard across the existing rate laws) is sufficient to guarantee the uniqueness of the steady state. The curious case of synergistic or mass-action rate laws has already been well-discussed by [29, 28] through constructing a particular Lyapunov function obtained from linear programming. Therefore, the analysis with Assumptions (6) and (7) brings completeness to the study of the global stability of biological networks.

Finally, contrary to the conclusions drawn through the linear analysis, the proposed methodology reveals the inability of the feed-forward loop to retain the global adaptive property. The linearised analysis leads to a system of linear algebraic equations with a trivial solution for the (linearised) downstream system. In contrast, due to the nonlinearity of the rate functions, the actual steady-state solutions for the downstream states are likely to be non-zero and dependent on the disturbance input thereby resulting in the violation of the adaptation property of the upstream feed-forward module.

Moreover, the linearised or local analysis of the system reveals the necessary paths (with its sign) and loops for perfect adaptation. In contrast, the practical constraints and global stability conditions allow us to zoom in further to obtain the appropriate edge configurations out of all the candidate loops and paths capable of local, perfect adaptation. Subsequently, investigating the robustness of each network structure admissible for global adaptation and obtaining appropriate structural prediction for robust, perfect, and global adaptation can be an exciting area of further study.

## Acknowledgements

PB acknowledges funding from the Ministry of Human Resources, India, and centre for Integrative Biology and Systems (m)Edicine (IBSE), IIT Madras.

## 5. Data Availability

The MATLAB codes can be accessed at https://github.com/priyan295/Nonlinear_Adap

