## Supplementary Material for "Design principles for perfect adaptation in biological networks with nonlinear dynamics"

### Supporting Information

#### 1. Proof for proposition 1

*Proof.* The trivial scenario of feed-forward structure can be safely ruled out as the system's steady state possesses a functional dependence on the external input. Therefore the remaining structural option for the two-node network is a feedback topology.

We shall prove through contradiction that it is necessary for the network to contain a buffer module over and above a feedback architecture. We first start with a two-node network where the concentration of the input-receiving node ($\mathcal{A}$) is denoted as $x_1(t)$ and the same for the other node ($C$) is $x_2(t)$. Given $x_2(t)$ is the output node, the corresponding dynamical system can be written as

$$\begin{aligned} \dot{v} &= f_v(v) \\ \dot{x}_1 &= f_{11}(x_1, x_1) + f_{12}(x_1, x_2) + f_{1v}(x_1, v) \\ \dot{x}_2 &= f_{21}(x_2, x_1) + f_{22}(x_2, x_2) \\ y(t) &= x_2(t) \end{aligned}$$

Since the present work focuses on constant (step or staircase type) disturbances, we shall limit our discussion to the case $f_v(v) = 0$.

$$f_{11}(x_1(v), x_1(v)) + f_{12}(x_1(v), x_2(v)) + f_{1v}(x_1(v), v) = 0 \quad (1)$$

$$f_{21}(x_2(v), x_1(v)) + f_{22}(x_2(v), x_2(v)) = 0 \quad (2)$$

Again, according to the condition in (12), replacing the output state $x_2(v)$ with a constant ($k_0$) we obtain

$$f_{11}(x_1(v), x_1(v)) + f_{12}(x_1(v), k_0) + f_{1v}(x_1(v), v) = 0 \quad (3)$$

$$f_{21}(k_0, x_1(v)) + f_{22}(k_0, k_0) = 0 \quad (4)$$

$$\implies x_1(v) = g(k_0) \quad (5)$$

$$\implies k_0 = \zeta(v) \text{ (Contradiction)} \quad (6)$$

As it can be inferred from (3) the only way to satisfy (11) and (12) is to not have any edge from the input-receiving node ($x_1$) to the output node ($x_2$), but it can be shown (Supporting information) that this shall render the system uncontrollable failing to satisfy the condition in (9) for perfect adaptation. Therefore, it can be concluded that a two-node network with an output node different from the input-receiving node cannot adapt.

Interestingly, if $x_1$ is considered as the output node *i. e.* $x_1(v) = k_1$ (constant), the infinite precision condition for adaptation can be written as

$$f_{11}(k_1, k_1) + f_{12}(k_1, x_2(v)) + f_{1v}(k_1, v) = 0 \tag{7}$$
$$\implies x_2(v) = g_1(v) \tag{8}$$
$$f_{21}(g_1(v), k_1) + f_{22}(g_1(v), g_1(v)) = 0 \tag{9}$$

Equation (9) can only be achieved if $f_2$ is made independent of $x_2$ which is possible for a class of rate kinetics prevalent in biochemical systems. Further, the corresponding condition for local stability requires the system matrix for the linearised system is Hurwitz which is only possible if node 1 ($x_1$) and 2 ($x_2$) engage in the negative cycle. □

### 2. Proof for proposition 3

It is worth mentioning that other than the aforementioned condition, equation (9) can also be satisfied if $f_2(.)$ can be factorized in the following manner

$$f_2(\mathbf{x}) = \hat{f}_2(x_1(v)) \times \tilde{f}_2(x_2) \tag{10}$$

where, $\hat{f}_2(x_1(v))$ has a singularity at $x_1 = k$.

However, equation (10) is not permitted under the confines defined by assumptions 1 – 6. Nevertheless, the existence of negative feedback would still be a necessity for local stability.

*Proof.* For possible scenario where, $\mathcal{A} \to \mathcal{B}$, $\mathcal{B} \to C$, $\mathcal{A} \to C$, the adaptation equations can be written as

$$f_{11}(x_1(v), x_1(v)) + f_{1v}(x_1(v), v) = 0$$
$$\implies x_1(v) = g_1(v)$$
$$f_{21}(k_0, x_1(v)) + f_{22}(k_0, x_2(v)) + f_{23}(k_0, x_3(v)) = 0$$
$$f_{31}(x_3(v), k_0) + f_{33}(x_3(v), x_3(v)) = 0$$
$$\implies x_3(v) = g_3(x_1(v))$$

Let the change in the disturbance level is $v_1$ to $v_2$. Due to the change in the disturbance level, the change in $f_{ij}$ be $\Delta f_{ij}$. The modified condition for adaptation can be written as

$$\Delta f_{23}(\Delta x_3) + \Delta f_{21}(x_1, \Delta x_1) = 0 \tag{11}$$
$$\implies \Delta f_{23} \circ \Delta g_3(x_1, \Delta x_1) + \Delta f_{21}(x_1, \Delta x_1) = 0 \tag{12}$$

where '$\circ$' denotes the composition of two functions.

It can be shown that due to the class $\mathcal{K}$ nature of $|f_{ij}|$ with respect to $x_j$, $x_1$, for this structure, possesses a monotonic relationship with $v$. Therefore, if $\Delta v := v_2 - v_1 > 0 \implies \Delta \geq 0 \implies \Delta|f_{21}| > 0$ owing to the class $\mathcal{K}$ nature of $f_{21}$. Therefore, the only way to satisfy equation (12) is to have a *mutual opposition* between the edge from $\mathcal{A}$ to $C$ and the forward path $\mathcal{A} \to \mathcal{B} \to C$. □

### 3. Proof for Theorem 1

*Proof.* Consider an $N-$node network with each node species' concentration as the state variable. Without any loss of generality, let us denote the concentration of the input-receiving node as $x_1(t)$.

We shall attempt to prove this by contradiction, *i.e,* hypothesize that a feed-forward $N-$node network with $< N$ edges can provide adaptation. It is trivial that an $N-$node network is structurally controllable only if it contains $\geq N - 1$ links [35]. The resultant network structure, in this case, cannot contain any loops to ensure structural controllability. Further, each node must be connected to the input-receiving node by one forward link. Therefore, the resultant graph shall always be isomorphic to a *hub and spoke* network, with the hub being the input-receiving node. Furthermore, there exists no path between the nodes of two different spokes. Suppose the output node is situated at the $k^{\text{th}}$ spot of $p^{\text{th}}$ spoke. Therefore, the perfect adaptation condition for the nodes in that branch.

$$\dot{v} = 0 \tag{13}$$

$$\begin{bmatrix} \dot{x}_1 \\ \dot{x}_2 \\ \vdots \\ \dot{x}_k \\ \dot{x}_{k+1} \\ \vdots \end{bmatrix}_{\mathbf{x}=\mathbf{x}^*} = \begin{bmatrix} f_{11}(x_1) + f_{1v}(x_1, v) \\ f_{21}(x_2, x_1) + f_{22}(x_2) \\ \vdots \\ f_{kk-1}(x_k, x_{k-1}) + f_{kk}(x_k) \\ f_{k+1k}(x_{k+1}x_k) + f_{k+1k+1}(x_{k+1}) \\ \vdots \end{bmatrix}_{\mathbf{x}=\mathbf{x}^*} = \mathbf{0} \tag{14}$$

Further, putting $x_k^* = k_0$ (constant) for infinite precision, we obtain from Eq. (14) $x_k^* = k_0 \implies x_1 = k_1 \implies v = f(k_1) = k_2$ where $k_0$, $k_1$ and $k_2$ are constants. This contradicts the assumption since $v$ is an external variable subject to fluctuation. Therefore, a network with $N$ nodes requires at least $N$ edges to perform global, perfect adaptation. □

### 4. Retroactivity

This section expands on the section about the effect of downstream connections on the adaptation-capable modules.

*4.1. Modularity of Negative feedback with buffer modules*

The dynamical system underlying an NFB network (in Equations (25) – (27)) can be expressed at steady as

$$
\begin{aligned}
f_{N,N-1}(x^*_{N-1}) - k &= 0 && (15)\\
\implies x^*_{N-1} &= k_1 && (16)\\
f_{N-1,N-2}(x^*_{N-1}, x^*_{N-2}) - f_{N-1}(x^*_{N-1}) &= 0 && (17)\\
\implies x^*_{N-2} &= k_2 && (18)\\
& && (19)
\end{aligned}
$$

Suppose the output node is the $K^{\text{th}}$ node. The $(K+1)^{\text{th}}$ steady state equation can be written as

$$
\begin{aligned}
f_{K+1,K}(x^*_{K+1}, x^*_K) - f_{K+1}(x^*_{K+1}) = 0 && (20)\\
\implies x^*_K = \tilde{k} \quad (x^*_{K+1} \text{ is constant due to previous equation.}) && (21)
\end{aligned}
$$

Therefore, as long as the stability is preserved, the modularity for negative feedback with buffer modules remains unaffected by downstream connections.

*4.2. Modularity of Incoherent feedforward modules*

Let us consider an adaptation-capable, feed-forward network along with a downstream node $x_d$ and the output being the concentration of $(j+1)^{\text{th}}$ node.

$$
\begin{aligned}
\dot{v} &= 0 && (22)\\
\dot{x}_1 &= f_{1,v}(x_1, v) - f_{1,1}(x_1) && (23)\\
\dot{x}_j &= \sum_{k=1}^{j-1} f_{j,k}(x_j, x_k) - f_{j,j}(x_j) && (24)\\
\dot{x}_{j+1} &= \sum_{k=1}^{j} f_{j+1,k}(x_j, x_k) - f_{j+1,j+1}(x_{j+1}) \pm f_{j+1,d}(x_{j+1}, x_d) && (25)
\end{aligned}
$$

The condition for perfect adaptation in the isolated scenario is

$$
\Delta \sum_{k=1}^{j} f_{j+1,k}(x_j, x_k) = 0
$$

where 'Δ' refers to the change in the functional values due to the change in the disturbance level $v$. Further, the condition for the same in the presence of the downstream connection can be written as

$$\Delta f_{j+1,d}(x_{j+1}, x_d) = 0 \tag{26}$$

Evidently, Equation (26) puts the baggage of producing adaptation to the downstream state $x_d$, showcasing the inability of feed-forward networks to provide global, perfect adaptation in the presence of downstream connection. However, Equation (26) can be satisfied if the steady state of $x_d$ is kept to zero. In that case, due to the introduction of negative feedback in the otherwise feed-forward structure, the corresponding response for $x_d$ shall be oscillatory, leading to a 'negative,' value thus an infeasible value of concentration.
